# Maternal dietary deficiencies in one-carbon metabolism during early neurodevelopment result in larger damage volume, reduced neurodegeneration and neuroinflammation and changes in choline metabolites after ischemic stroke in middle-aged offspring

**DOI:** 10.1101/2023.02.15.528759

**Authors:** Lauren Hurley, Jesse Jauhal, Sharadyn Ille, Kasey Pull, Olga V. Malysheva, Nafisa M. Jadavji

## Abstract

Maternal diet pregnancy and lactation is vital to the early life neuro programming of offspring. One-carbon (1C) metabolism, which includes folic acid and choline, plays a vital role in closure of the neural tube and other neurodevelopment. However, the impact of maternal dietary deficiencies on offspring neurological function following ischemic stroke later in life remains undefined. Stroke is one of the leading causes of death globally, and its prevalence is expected to increase in younger age groups as the incidence of various risk factors for stroke increases. Furthermore, our group has shown that dietary deficiencies in 1C metabolism result in worse stroke outcome. Our study aimed to investigate the role of maternal dietary deficiencies in folic acid and choline on ischemic stroke outcome in middle-aged male and female mice. Female mice were maintained on either a control or deficient diets prior to and during pregnancy and lactation. When female and male offspring were 10-months of age, ischemic stroke was induced via photothrombosis targeting the sensorimotor cortex. Stroke outcome was assessed by measuring motor function in living animals and ischemic damage volume, neurodegeneration, neuroinflammation, and choline metabolism in the brain postmortem. No significant difference was observed between maternal dietary groups in offspring motor function; however, males and females differed in their motor function. Maternal diet significantly impacted ischemic damage volume. Male and female offspring from deficient mothers showed significantly reduced neurodegeneration and neuroinflammation within the ischemic damage region. We also report changes in plasma 1C metabolites as a result of maternal diet and sex after ischemic stroke in offspring. Our data indicates that maternal dietary deficiencies do not impact offspring motor outcome following ischemic stroke, but do play a role in other ischemic stroke outcomes such as ischemic damage volume and plasma 1C metabolites in middle-aged adult offspring. Furthermore, our data indicates that sex of mice plays an important role in stroke outcome during middle-age.

## 1. Introduction

Stroke prevalence in young adults and the underlying causes are of increasing concern [1–6]. In Brazil ischemic stroke incidence was found to increase in adults less than 45 years-old from 2005 to 2011 by 88% and by 62% from 2005 to 2015 [2]. Rates of hospitalizations for ischemic stroke in young adults have been increasing in European countries since 1987 including Denmark [6], France [1,7], and Sweden [4]. In the Greater Cincinnati/ Northern Kentucky region of the United States, proportion of stroke for people under 55 years-old increased from 12.9% in 1993 to 18.6% in 2005 [8] and the U.S. as a whole saw an increase in age-specific hospitalization rates for acute ischemic stroke in patients aged 25 to 44 years and 45 to 64 years-old [9].

Nutrition is a modifiable risk factor for stroke [10–12]. B-vitamins, such folic acid, or folate, as well as choline are known for their important role in neurodevelopment [13–15]. Both folic acid and choline play a major role in one-carbon (1C) metabolism, which is responsible for methylation metabolism, maintenance of cellular redox status, nucleotide metabolism, and lipid biosynthesis [16,17]. These vitamins have also been implicated in the onset and progression of stroke [18–20], as insufficient folic acid and choline levels during early life development are shown to have increased risk of stroke and cardiovascular disease in humans and animals [21].

During pregnancy, there is an increased demand for dietary nutrients and increased cognitive performance throughout all stages of life has been seen when there is adequate levels of choline in maternal nutrition [22]. Maternal deficiencies in dietary folic acid and choline result in increased homocysteine, which hinders neurodevelopment, such as hippocampal and cerebellum development, and impacts short-term memory function in offspring [14,23]. Alterations in the 1C metabolic pathway due to dietary deficiencies like dietary choline and folate can also cause increased oxidative stress and cell death in the hippocampus of offspring [14,23,24], leading to improper neurodevelopment and profound negative impacts both in the short- and long-term. Inadequate levels of choline and folate during pregnancy can also decrease cognitive abilities of offspring later in life, including increased cognitive decline and neurodegeneration [22]. Overall, maternal nutrition contributes to early life programming and development of the fetus, with lasting effects in both early and late life of offspring [25]. The aim of this study was to investigate the role of maternal dietary deficiencies in folic acid and choline on ischemic stroke outcome in middle-aged offspring.

## 2. Materials and Methods

All experiments involving the use of animals have been approved by the Midwestern University IACUC (2983). Experimental manipulations are summarized in Figure 1. Maternal females were born and raised on a control diet until two months of age, then were placed on either a folic acid (FADD) or choline deficient diet (ChDD) or control diet (CD). Both male and female parent mice were placed on these diets for 4 weeks prior to mating. Female mice were maintained on either a control diet or deficient diet during pregnancy and lactation. Female and male offspring were weaned and placed on a control diet until ten months of age then had ischemic stroke induced via photothrombosis targeting the sensorimotor complex. Four weeks after ischemic damage, the offspring underwent behavioral testing to assess motor function and after completion were euthanized with blood and brain tissue samples collected for analysis.

**Figure 1:**
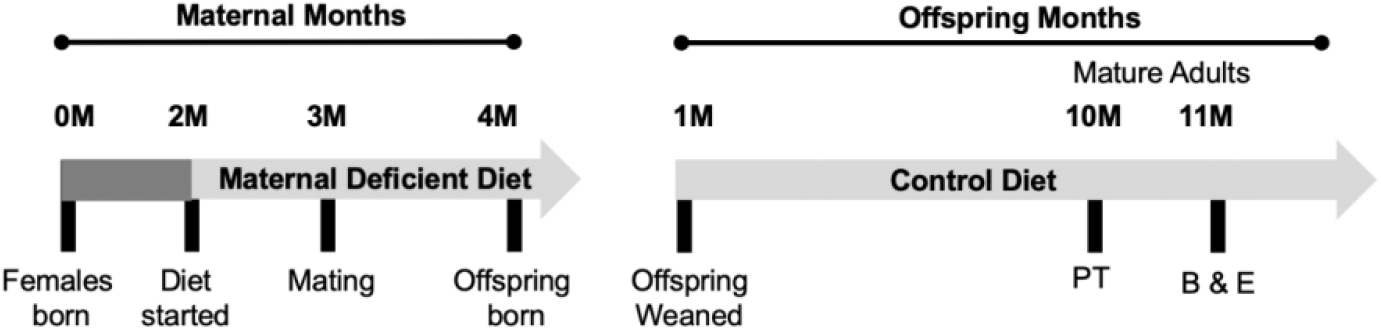
Summary of experimental manipulations. Maternal mice are born and raised with a conventional diet until two months of age, after which they are assigned experimental diets. Females are mated with males of the same corresponding diet at three months and offspring are born one month after mating. After offspring are weaned, they are placed on a control diet that is continued until ten months of age, at which point the mice underwent photothrombosis. After one month of recovery, the offspring undergo behavioral testing and then are euthanized, and tissue samples collected. Maternal months consists of the time from which the female maternal mice are born until the birth of offspring. Offspring months consist of the time from which offspring area weaned until they were euthanized and tissue samples were collected. Abbreviations, behavior (B), euthanization (E) and photothrombosis (PT).

### 2.2. Dietary Composition

The maternal diets in this study consisted of a control diet (CD), folic acid deficient diet (FADD), and choline deficient diet (ChDD). The CD represented normal levels of both folic acid and choline that would be found in a conventional diet, which were determined from previous literature and experimentation[23,26,27], both mothers and offspring were fed these diets. The deficient diets consisted of lower levels of folic acid or choline compared to the control diet. The CD contained 2 mg/kg of folic acid and 1,150 mg/kg of choline bitrate, whereas the FADD contained only 0.3 mg/kg and the ChDD only had 300 mg/kg of choline bitrate.

### 2.3. Photothrombosis Model

The photothrombosis model is a reproducible model that induces ischemic damage by targeting the sensorimotor cortex [28]. The mice were anesthetized in an isoflurane chamber and the surgical site was prepared by shaving and disinfecting the head.

Anesthesia was continued via face mask and body temperature was maintained at 37±C with a heating pad throughout the entire procedure. To create ischemic damage and simulate stroke, Rose Bengal, a photoactive dye, was injected intraperitoneally at a dose of 10 mg/kg five minutes prior to laser exposure. Activation of the dye with a 532nm laser beam centered 3 cm over the sensorimotor cortex for 15 minutes caused formation of free radicals to induce endothelial injury, platelet activation and aggregation resulting in ischemic damage.

### 2.4. Behavioral Testing

Behavioral testing in the offspring was conducted four weeks after ischemic damage was induced via photothrombosis to assess motor function. This was done using accelerating rotarod, forepaw placement, and ladder beam walking tasks.

#### 2.4. a. Accelerating Rotarod

Coordination and balance were tested by placing mice on an accelerating rotarod and measuring latency to fall and rotations per minute. This was done for three trials per mouse with five-minute lapse periods between trial.

#### 2.4. b. Forepaw Placement

The forepaw placement task assesses forepaw usage by video recording mice placed in a glass cylinder (19cm high, 14cm diameter) for ten minutes and analysing contact between the walls of the cylinder and forepaw of the mouse [28]. The first 20 rears were analyzed during this time.

#### 2.4. c. Ladder Beam Walking

Skilled motor function of impaired or non-impaired fore- and hindlimbs was measured by ladder beam walking. Mice were placed on a horizontal ladder composed of two plexi-glass walls with metal bars inserted into the holes at the bottom edge of the walls. The mice were assessed in their ability to travel from one end towards the other, without being permitted to regress. After being familiarized with the ladder, the remaining trials were video-recorded and performance was analyzed in each frame for quality of limb placement as previously described [29].

### 2.5. Brain Tissue Collection and Sectioning

After completing behavioral analysis and euthanasia, brain tissue was collected and sectioned using a cryostat. Tissue sections were 30 μm thick and mounted onto positively charged microscope slides. Serial sections were collected for each brain tissue sample over six slides total. A minimum of 8 sections within the damaged region of the brain were collected. Slides were stored −80±C until tissue analysis was conducted. For damage volume analysis, each animal had a minimum of three sections of damaged area per slide and two slides per animal were stained using cresyl violet (Sigma). Images were taken using an Olympus microscope. ImageJ (NIH) software was used to quantify volume of damaged tissue by measuring the area of damage in each serial section [30] and multiplying the area between sections (0.03mm) by the number of sections from the damaged site.

### 2.6. Immunofluorescence Staining

To investigate neurodegeneration and neuroinflammation, immunofluorescence analysis of ischemic-damaged brain tissue was performed. To assess neurodegeneration by measuring apoptosis in ischemic-damaged brain tissue, primary antibodies to active capase-3 (1:100, Cell Signaling Technologies) were used. All brain sections were stained with a marker for neuronal nuclei, NeuN (1:200, AbCam). Primary antibodies were diluted in 0.5% Triton X and incubated on the brain tissue overnight at 4°C. Brain sections were incubated with secondary antibodies Alexa Fluor 488 or 555 (1:200, Cell Signaling Technologies) the following day at room temperature for 2 hours then stained with 4’, 6-diamidino-2phenylindole (DAPI, 1:10000).

To assess neuroinflammation by measuring active microglial cells, primary antibodies to Ionized calcium binding adaptor molecule 1 (Iba1, 1:100, AbCam) and CD68 (1:500, Bio-Rad) were used. Primary antibodies were diluted in the same manor and incubated on the brain tissue overnight at 4°C. Anti-rat (594) and anti-rabbit (488, Cell Signaling Technologies) secondary antibodies were added the following day and incubated at room temperature for 2 hours then stained with DAPI (1:10000).

Microscope slides were coverslipped with fluromount and stored at 4°C until analysis. The staining was visualized using an ECHO microscope and all images were collected at the magnification of 200X. To assess neurodegeneration within the ischemic core region of the brain tissue, co-loacalization of active caspase-3 with NeuN and DAPI labelled neurons were counted and averaged per animal. To assess neuroinflammation, co-localization of Iba1 and CD68 labelled cells were counted. Cells were distinguished from debris by identifying a clear cell shape and intact nuclei (indicated by DAPI and NeuN) under the microscope. All cell counts were conducted by two individuals blinded to treatment groups. Using ImageJ, the number of positive cells were counted in three brain sections per animal, a total of 5 animals were analyzed per group. For each section, three fields were analyzed. The number of positive cells were averaged for each animal.

### 2.8. Plasma One-Carbon Metabolite Measurements

Plasma tissue from offspring was measured for betaine, choline, dimethylglycine, methionine, phosphocholine, phosphatidylcholine, lysophosphatidylcholine, and, Trimethylamine N-oxide (TAMO), sphingomyelin levels using the LC-MS/MS (ThermoElectron Corp, Waltham, MA) method as previously reported [31–33].

### 2.9. Data Analysis and Statistics

GraphPad Prism 6.0 was used to analyze behavioral testing, plasma tHcy measurements, lesion volume, immunofluorescence staining, and plasma choline measurements. In GraphPad Prism 6.0, two-way ANOVA analysis was performed when comparing the mean measurement of both sex and dietary group for behavioral testing, one-carbon metabolites, lesion volume, and immunofluorescence staining. Significant main effects of two-way ANOVAs were followed up with Tukey’s post-hoc test to adjust for multiple comparisons. All data are presented as mean + standard error of the mean (SEM). Statistical tests were performed using a significance level of 0.05. Behavioral and brain tissue microscope data were analyzed by two individuals that were blinded to experimental treatment groups.

## 3. Results

### 3.1. Accelerating Rotarod

The accelerating rotarod was used to assess coordination and balance by measuring the length of time spent on the rotarod, termed latency to fall. No significant difference was noted between maternal diet groups (Figure 2A; F(_2,37_) = 1.25, p = 0.30) and no interaction was observed between maternal diet and sex main effects (Figure 2A; F(_2,37_) = 0.60, p = 0.56). Significant sex differences between male and female offspring were noted with performance (Figure 2; F(_1,37_) = 18.04, p = 0.0001). No significant difference was observed between maximum speed (rotations per minute, RPM) achieved between groups (F(_2,37_) = 1.78, p = 0.18), however female mice reached a higher RPM compared to males (F(_2,37_) = 21.92, p < 0.0001). No significant interaction between maternal diet and sex was seen (F(_2,37_) = 0.49, p = 0.62).

**Figure 2:**
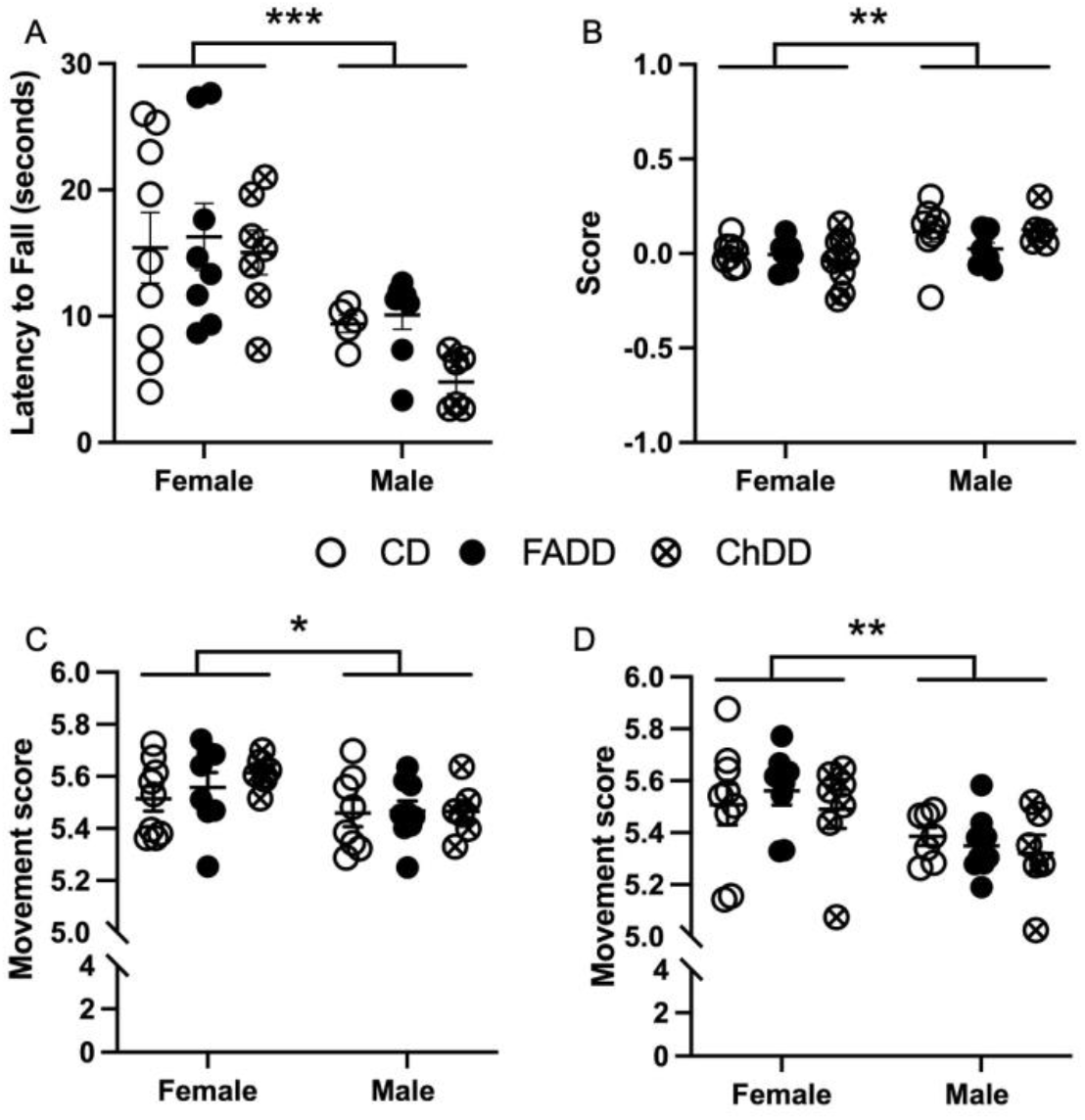
Impact of maternal diet on skilled motor function in 11-month-old male and female mice one-month after induction of ischemic stroke. (**A**) Latency to fall on acclerating rotarod. (**B**) Score on forepaw placement task. Movement score on ladder beam walking task (**C**) impaired and (**D**) non-impaired forepaw. Depicted are means of + SEM of 10 to 11 mice per group. Two-way ANOVA statistical analysis was performed on data. * p < 0.05, ** p < 0.01, ** p < 0.001 main sex difference, 2-way ANOVA.

### 3.2 Forepaw placement

Impaired forepaw usage during forepaw placement task was not significantly impacted by maternal diet (F(_2,41_) = 0.33, p = 0.72) or sex (F(_1,41_) = 0.83, p = 0.37) and no significant interaction was observed between sex and maternal diet for impaired (F(_2,41_) = 0.44, p = 0.64) or non-impaired forepaw usage (F(_2,41_) = 0.05, p = 0.95). Non-impaired forepaw usage also did not have a significant main effect of maternal diet (F(_2,41_) = 0.05, p = 0.95) or sex (F(_1,41_) = 3.01, p = 0.09), or interaction between sex and maternal diet (F(_2,41_) = 0.77, p = 0.47). No main effect of maternal diet (F(_2,40_) = 0.78, p = 0.46) was seen with forepaw placement score, which is the ratio of impaired to non-impaired forepaw usage, however a significant main effect of sex was observed (Figure 2B; F(_1,40_) = 9.98, p = 0.003).

### 3.3 Ladder Beam

The ladder beam test was used to measur skilled motor function. The test consisted of a movement score based on usage of impaired and non-impaired fore- and hindlimb stepping patterns, as well as the number of errors made and time to cross the ladder. For impaired movement score, there was no difference in maternal dietary groups (Figure 2C; F(_2,41_) = 0.61, p = 0.55), but a sex difference was present (Figure 2C; F(_1,41_) = 6.61, p = 0.01) and no interaction between sex and maternal diet (F(_2,41_) = 0.48, p = 0.62). The movement score for non-impaired fore and hindlimbs showed no difference in maternal dietary groups (Figure 2D; F(_2,40_) = 0.35, p = 0.71), but a sex difference was present (Figure 2D; F(_1,41_) = 11.04, p = 0.01) and no interaction between sex and maternal diet was present (F(_2,40_) = 0.28, p = 0.75).

Percent error, number of errors made while crossing the ladder beam, showed no differences between maternal diet for impaired (F(_2, 41_) = 1.31, p = 0.28) and non-impaired limbs (F(_2, 42_) = 0.03, p = 0.97). There was also no sex difference in the number of errors made for impaired (F(_1, 41_) = 3.6, p = 0.06) and non-impaired limbs (F(_1, 41_) = 1.04, p = 0.31). No interaction between sex or maternal diet with impaired percent error (F(_2,41_) = 0.23, p = 0.80) or non-impaired percent error (F(_2,42_) = 0.75, p = 0.48) were observed. Time to cross the ladder beam was not significantly impacted by maternal diet (F(_2,42_) = 1.22, p = 0.31) or sex (F(_1,42_) = 0.14, p = 0.11), there was also no interaction sex and maternal diet (F(_2, 42_) = 0.15, p = 0.86).

### 3.4 Ischemic damage volume increased in offspring as a result of maternal dietary deficiencies

Brain tissue sections stained with cresyl violet (Sigma) and imaged using an Olympus microscope (Figure 3A). Images were processed using ImageJ software to measure ischemic damage volume. Maternal diet had a significant effect on damage volume (Figure 3B; F(_2, 32_) = 3.53, p = 0.041), but no main effect of sex (Figure 3B; F(_1,32_) = 0.63, p = 0.43) or significant interaction between maternal diet and sex for ischemic damage volume was observed (Figure 3B; F(_2,32_) = 0.24, p = 0.79).

**Figure 3:**
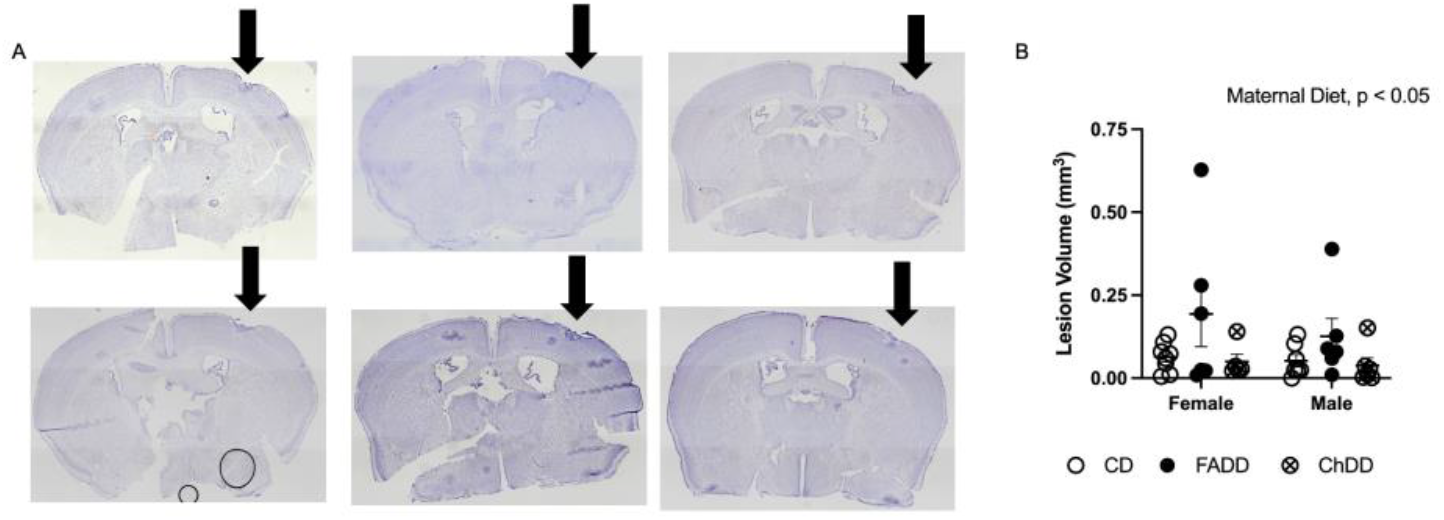
Cresyl violet-stained brain tissue sections and ischemic damage volume in offspring. Impact of maternal diet on ischemic damage volume after stroke in 10-month-old male and female mice. (A) Images depicting ischemic-damaged region of brain tissue after stroke in male and female mice of each maternal dietary group; damage indicated by blue arrow. (B) Quantification of lesion volume (mm^3^). Depicted are means + SEM of 6-9 mice per sex and diet group.

### 3.5 Decreased neurodegeneration and neuroinflammation in male and female offspring from deficient mothers

Within the ischemic damaged region of the brain tissue, we measured neurodegeneration and neuroinflammation. Neurodegeneration was assessed using active caspase-3 levels, representative images are shown in Figure 4A. Maternal diet had an impact on the number of positive active-caspase 3 neurons (Figure 4B; F(_2, 13_) = 31.22, p<0.001). There was no effect of sex (F (_1, 13_) = 0.53, p = 0.48) and no interaction between sex and maternal diet (F (_2, 13_) = 0.18, p = 0.84).

**Figure 4.**
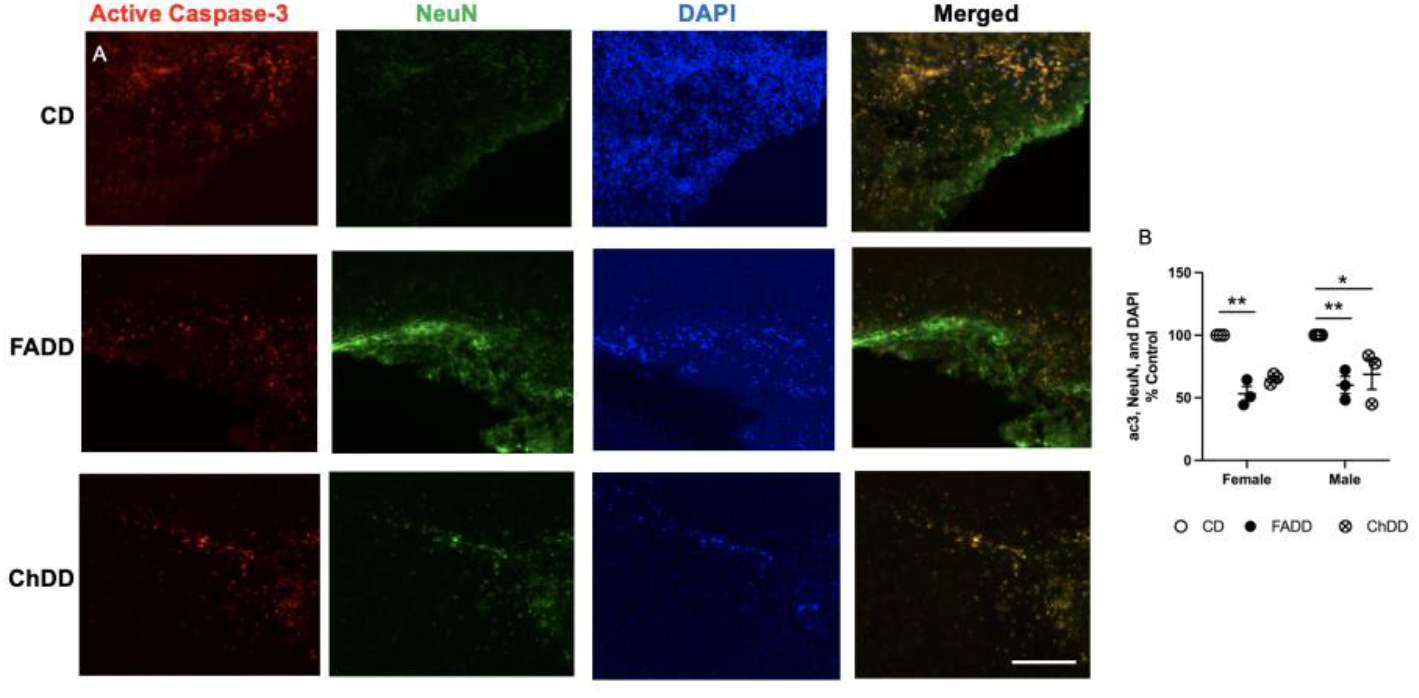
Impact of maternal dietary deficiencies on offspring neurodegeneration (neuronal apoptosis) in an ischemic brain region 1.5 months after photothrombosis surgery. (A) Representative images for active caspase-3, neuronal nuclei (NeuN) and 4’,6-diamidino-2-phenylindole (DAPI) staining and quantification of active caspase-3, NeuN and DAPI cell counts. (B) Scale bars: 50 μm. Depicted are means of + SEM of 3 to 4 mice per group. * p < 0.05, ** p < 0.01, maternal diet main effect. Abbreviations: CD, control diet; ChDD, choline deficient diet and FADD, folic acid deficient diet.

We measured neuroinflammation within the ischemic damage region using Iba1 and CD68 colocalization [34], representative images are shown in Figure 5A. Maternal diet had an impact on the number of positive active-caspase 3 neurons (Figure 5B; F (_2, 22_) = 5.51, p=0.01). There was no effect of sex (F (_2, 22_) = 2.82, p = 0.11) and no interaction between sex and maternal diet (F (_2, 22_) = 2.49, p = 0.11).

**Figure 5.**
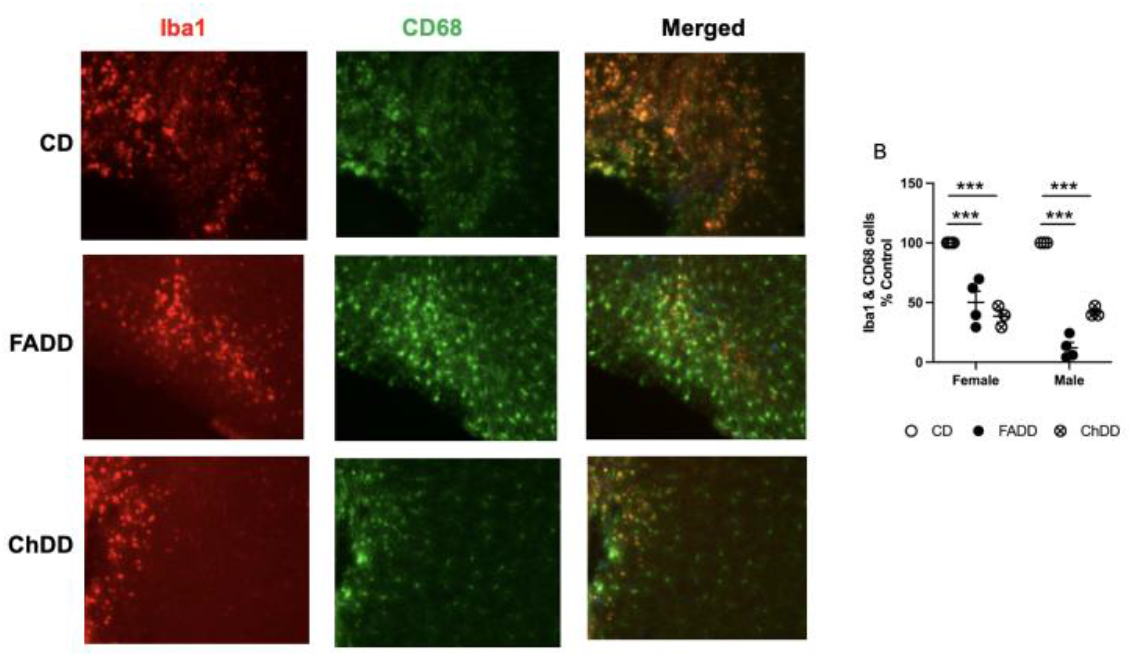
Impact of maternal dietary deficiencies on offspring neuroinflammation in ischemic brain region 1.5 months after photothrombosis surgery. (A) Representative images for Iba1 and CD68 staining. (B) Quantification of Iba1 and CD68 cell counts. Scale bars: 50 μm. Depicted are means of + SEM of 3 to 4 mice per group. *** p < 0.001, maternal diet main effect. Abbreviations: CD, control diet; ChDD, choline deficient diet and FADD, folic acid deficient diet.

### 3.6 Plasma one-carbon metabolites

We measured one-carbon metabolites in plasma of offspring 1.5 months after ischemic stroke and found that maternal diet impacted methionine (Figure 6: F (_2, 43_) = 3.24, p 0.049), choline (F (_2, 43_) = 6.65, p=0.003), DMG (F (_2, 43_) = 4.60, p = 0.0001), PC (F (_2, 43_) = 5.86, p = 0.006), SM (F (_2, 43_) = 9.74, p = 0.003), and LPC (F (_2, 43_) = 14.91, p < 0.0001). There were pairwise differences between males from CD and FADD mothers for choline (p = 0.026), PC (p = 0.0032), and SM (p = 0.0011). There was also a difference between females and male animals for the following metabolites: choline (F (_1, 43_) = 5.15, p = 0.03), PC (F(_1, 43_) = 77.66, p < 0.001), SM (F (_1, 43_) = 48.08 p < 0.001), and LPC (F (_1, 43_) = 13.7, p = 0.0006). There was no sex difference in methionine (F (_1, 43_) = 1.96, p = 0.17) and DMG (F (_2, 43_) = 2.4, p = 0.11). A significant interaction between maternal diet and sex were found for PC (F (_2, 43_) = 3.02, p = 0.06), no other metabolites had an interaction (methionine, F (_2, 43_) = 0.071, p = 0.93; choline, F (_2, 43_) = 1.04. p = 0.36; DMG, Interaction: F (_2, 43_) = 2.35, p = 0.11; SM, F (_2, 43_) = 1.33, p = 0.28 and LPC, Interaction: F (_2, 43_) = 0.26, p = 0.77)

**Figure 6.**
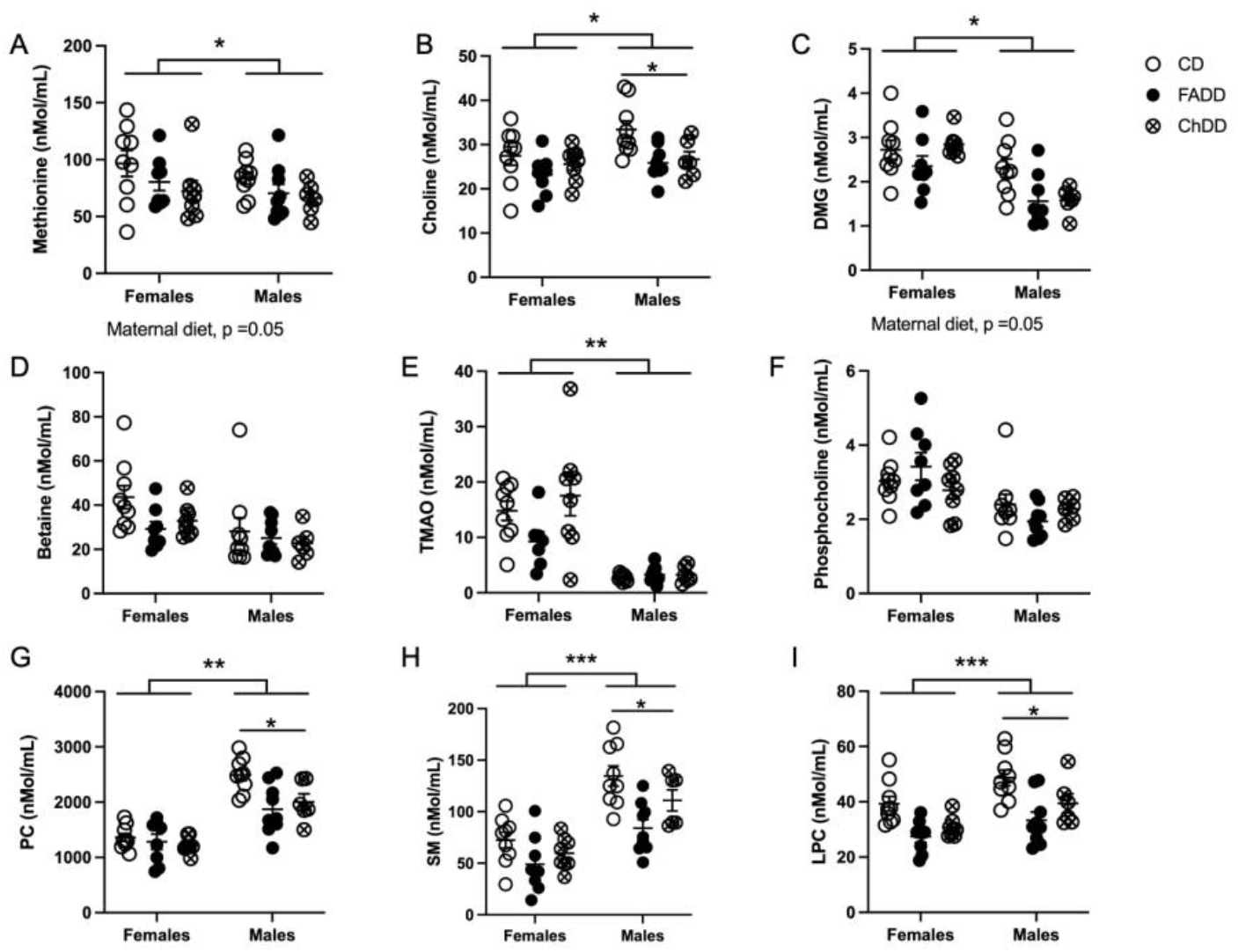
Impact of maternal dietary deficiencies on plasma choline metabolite levels 1.5 months after photothrombosis surgery. Levels of methoine (A), choline (B), dimethylglycine (DMG, C), betaine (D), Trimethylamine N-oxide (TAMO, E) phosphocholine (F), phosphatidylcholine (PC, sphingomyelin (SM, H) and lysophosphatidylcholine (LPC, I). Depicted are means of + SEM of 8 to 10 mice per group. * p < 0.001, maternal diet main effect or Tukey’s pairwise comparison. Abbreviations: CD, control diet; ChDD, choline deficient diet and FADD, folic acid deficient diet.

## 4. Discussion

Maternal nutrition and diet play an important role in the neurodevelopment of offspring. Alterations in this early life nutritional programming can have long term impacts on offspring health [35]. A maternal diet rich in required vitamins and nutrients has a large impact in the prevention of various diseases including stroke [36]. What remains unclear is the impact of maternal nutritional in one-cardeficiencies during pregnancy and lactation, and their impact on offspring stroke outcome later in life. The aim of this study was to determine the impact of maternal deficiencies in folic acid and choline on middle-aged offspring stroke outcome. After ischemic stroke, no differences were seen between maternal dietary groups in the accelerating rotarod, forepaw placement, and ladder beam tasks. Sex differences were seen between females and males in the accelerating rotarod, forepaw placement, and ladder beam tasks indicating males had worse outcome. Analysis of brain tissue indicated that there was a significant main effect of maternal diet on ischemic damage volume, neurodegeneration, and neuroinflammation. Changes in plasma choline metabolites were observed as a result of maternal diet and sex after ischemic stroke in offspring.

Sex has also been correlated with stroke incidence in young adults, however which sex having the highest incident rate is still being debated. In the same Brazil study previously mentioned, ischemic stroke incidence was found to be higher in men less than 45 years-old and higher in women less than 55 years-old [2]. According to the Danish National Patient Register, when reviewing trends in young adults aged 15 to 30 years-old, incidence rates for hospitalization for ischemic stroke and transient ischemic attack (TIA) were high in women compared to men [6]. In France, ischemic stroke incidence significantly increased in men aged 25 to 74 years-old and in women aged 35 to 64 years-old from 2008 to 2014 [7]. In the same study, women with ischemic stroke were also found to have a higher mortality when between the ages of 45 and 64 years-old and men had significantly younger mean age at hospitalization [7]. Eastern France had a higher increase in incidence rates for ischemic stroke in men after 2003 as compared to before 2003 [1] and the U.S. had similar trends with age-adjusted acute ischemic stroke hospitalization rates being higher in men from 2000 to 2010 [37]. China, however, saw onset of first-ever stroke in significantly younger women compared to men, with the average age being 36.9 years-old compared to 38.7 years-old in men from 2007-2018 [5]. The high variation between regions could be due to the prevalence of risk factors associated with ischemic stroke in each sex.

In the present study there is no impact of maternal diet on motor function after ischemic stroke. We do describe a significant sex difference in behavioral outcome, females were observed to perform better on the motor function behavioral tests. These findings may be due to a variety of mechanisms, however the most plausible appears to be due to the neuroprotective effects of estrogen in female mice post-ischemic stroke [38]. The neuroprotective effects of estrogen on stroke are extremely well documented throughout the literature, with various studies demonstrating significantly greater damage post ischemia in male mice and rats when compared to females [39,40]. This was further confirmed through the removal of ovaries in female mice and rats, who subsequently appeared to have lost this protective effect after ischemia [41,42]. Therefore, the evidence for estrogen mediated neuroprotection after ischemic damage is quite clear and is likely the mechanism through which we observed the sex difference in this study. The estrous cycle of our female mice was not monitored, which could also provide an explanation for the variability seen within the females. As circulating concentrations of estrogen vary depending on the point at which a female is in in the estrous cycle, it is highly likely that females with high circulating estrogen during and after ischemic stroke induction had greater recovery when compared to those with lower estrogen concentrations during those times [42].

The folic acid deficient diet appears to be driving the main effect of diet seen in the ischemic damage volume quantification and plasma one-carbon metabolites.. This variation seen in the folic acid deficient groups may be due to a variety of reasons. The complexity of the 1C metabolic pathway and the various compensatory mechanisms in place in this network may have impacted the ability of the tissue to recover [43]. Furthermore, the amount of diet consumed, as the mice were given food *ad libitum*, may have varied and impacted the amount of damaged tissue. Overall, a variety of potential influences may have resulted in the differences seen in damage volume. When compared to behavioral tests, the lesion volume quantification is likely more sensitive to alterations in 1C [44]. There is more compensation at a behavioral level due to the complexity of brain pathways and often times damage and injury can be masked in behavior [45]. We have previously reported that a deficient diet during pregnancy and lactation impairs motor function in 3-month-old offspring [27], however, in this study we show that these effects are only present in brain tissue of middle-aged (11 month old) offspring and that there is behavioral compensation that occurs after ischemic stroke.

## 5. Conclusions

This data provides insight on the complexity of neurodevelopmental pathways during early life, specifically in the 1C metabolic network, and how various levels of compensation may be in place to help combat detrimental effects of dietary deficiencies in early life. Using this study as a basis for further research, the possible neurological mechanisms that allow for this compensation to occur may be explored. Furthermore, this study adds to the large amount of current research that investigates the impact of sex on ischemic stroke outcome.

## Supplementary Materials

The following supporting information can be downloaded at: www.mdpi.com/xxx/s1, Figure S1: title; Table S1: title; Video S1: title.

## Author Contributions

Conceptualization, N.M.J.; methodology, O.V.M., N.M.J.; formal analysis, L.H., J.J., S.I., O.V.M., N.M.J.; investigation, L.H., J.J., S.I., O.V.M., N.M.J.; resources, N.M.J; data curation, L.H., J.J., S.I., O.V.M., N.M.J.; writing—original draft preparation, J.J., L.H., O.V.M, N.M.J..; writing—review and editing, L.H., J.J, O.V.M, N.M.J.,.; visualization, L.H., J.J.,.; supervision, N.M.J.; project administration, N.M.J.; funding acquisition, N.M.J. All authors have read and agreed to the published version of the manuscript

## Institutional Review Board Statement

The animal study protocol was approved by the Institutional Review Board of MIDWESTERN UNIVERSITY (2983, February 24, 2020).

## Informed Consent Statement

Not applicable.

## Data Availability Statement

Not applicable.

## Acknowledgments

Not applicable.

